# Smaller Genomes and Shorter Prophage Sequences Differentiate Clinical from Environmental Isolates of *Pseudomonas aeruginosa*

**DOI:** 10.64898/2025.12.04.691226

**Authors:** Robert W. Targ, Patrick Blankenberg, Zachary Flamholz, Julie D. Pourtois, Niaz Banaei, Elizabeth B. Burgener, Carlos E. Milla, Paul L. Bollyky

## Abstract

The opportunistic bacterial pathogen *Pseudomonas aeruginosa* can live in both environmental reservoirs as well as in the airways of people with cystic fibrosis (pwCF) and the evolutionary adaptations that enable *P. aeruginosa* to transition between these settings have clinical implications. Here, we compare the genome composition, prophage-associated contigs, mobile genetic elements, and phylogenetic source patterns across 146 full *P. aeruginosa* genomes sampled from the airways of pwCF, and 140 genomes from a variety of environmental sources (286 total genome isolates). We find that CF respiratory genomes are smaller and have higher GC-content than environmental genomes (nested F-test p=1.15*10^-11^; adjusted R^2^=0.832). We also find that CF respiratory genomes had fewer high-confidence full-prophage-labeled contigs (IRR=0.583, 95% CI 0.506-0.672) while the overall number of prophages does not differ between the groups. Approximately 85% of annotated prophage-associated genes have an unknown function. In contrast to prophages, the overall proportions of integrative elements and transposons are broadly similar between groups. CF respiratory and environmental isolates are mixed throughout the core-genome phylogeny and stochastic mapping do not support a preferred direction of change between source states, implying that pwCF acquire *P. aeruginosa* from environmental sources but also that clinical lineages are present in the environment. Overall, these results support differences in bacterial genome composition and prophage-associated contig length between CF respiratory and environmental isolates.

**Importance:** Understanding how *Pseudomonas aeruginosa* adapts from environmental reservoirs to the airways of people with cystic fibrosis is critical for understanding infection acquisition, persistence, and evolution. Our findings identify genome reduction and altered prophage content as distinguishing features of CF respiratory isolates, while revealing extensive phylogenetic overlap between clinical and environmental populations. These results highlight prophages as an understudied component of *P. aeruginosa* adaptation and provide insight into the evolutionary changes accompanying the transition from environmental organism to chronic human pathogen.

## Introduction

Bacteria move among habitats that impose different nutrient, competition, immune, and antimicrobial pressures. Adaptation across those settings can involve changes in both core-genome sequence and accessory DNA received via horizontal gene transfer. Mobile genetic elements (MGEs), pieces of DNA that can move around within a genome or between cells, are therefore potential records of ecological history and possible sources of new traits.^1,25,29^

*Pseudomonas aeruginosa* is a gram-negative bacterium found in soil, water, built environments, and the human body.^1,6,7^ It is also an opportunistic pathogen that causes severe infections, including chronic lung infections in the elderly, the chronically ill, and lung transplant recipients^3–5,73–76^ Infection can come either from the environment or from person-to-person transmission.^4,6–17^ Unlike many human pathogens involved in chronic lung infections, *P. aeruginosa* is not part of the normal flora of most healthy individuals, such that many infection events represent an evolutionary transition from an environmental habitat to the human host

One setting where *P. aeruginosa* infections are particularly devastating is in cystic fibrosis (CF), a genetic disease associated with mutations in the CFTR gene, a transmembrane chloride channel. While healthy humans typically resist chronic infection with *P. aeruginosa* infections, in people with Cystic Fibrosis (pwCF), defective CFTR function results in thick, dehydrated mucus and impaired mucociliary clearance that is a favorable environment for *P. aeruginosa* to establish chronic lung infections^77,78^. The airways of pwCF present a unique environment, with bacteria navigating a highly complex spatial structure^79,80^. In addition, Pa has been shown to adapt to the CF airway and exhibit changes in metabolism, antibiotic resistance, motility, and other virulence factors^81–83,98–100^. There is therefore great interest in understanding the selective pressures and genomic adaptations involved in *P. aeruginosa* colonization of the CF respiratory environment.

To facilitate transitions between environmental habitats and human hosts, *P. aeruginosa* can make use of its extensive phenotypic and genomic diversity. Infection in a human can select for genes that contribute to biofilm formation, antimicrobial resistance, and altered virulence, whereas environmental persistence can require responses to nutrient limitation and intermicrobial competition.^3,16,18–24^ A large and flexible pan-genome and accessory genome further helps *P. aeruginosa* adapt to that variability.^1,25,26^ The *P. aeruginosa* accessory genome, which is a set of genes that some strains have and others do not, includes genomic islands, integrative elements, plasmids, insertion sequences, and prophages.^1,16,27^ Living in a host can sometimes favor losing genes that cease to be useful, whereas more variable environmental settings may favor keeping a wider range of accessory genes.^1,4,26,71^

There is a growing awareness of the contributions that bacteriophages (phages), viruses that infect bacteria, make to the accessory genome. While lytic phages shape the bacterial genome through natural selection, lysogenic phages can alter the DNA of the bacteria they infect by integrating into the bacterial chromosome as a prophage or as an extra-chromosomal element. These phage sequences typically aid in phage replication and also encode defense, virulence, or other accessory functions.^2,29,34–36,96,97^ In *P. aeruginosa*, some of the experimentally-demonstrated effects of lysogenic phages include (but are not limited to) impacts on biofilm formation, defense against the eukaryotic immune system, antibiotic response, and immunity to future phage infection.^27,37–46,84–90^.

However, the contributions that prophages make to the adaptation of *P. aeruginosa* to CF airways are unclear. Well-controlled studies comparing clinical and environmental isolates are lacking. Only recently have the computational tools and prophage databases been sufficiently robust to facilitate these comparisons.

Here, we have asked how prophages vary between *P. aeruginosa* isolates collected in clinical versus environmental settings. To this end, we compare the genomes of *P. aeruginosa* from CF respiratory infections of pwCF with genomes from environmental sources. We look at bacterial genome size and GC content, prophage-associated contigs, differences in classes of MGEs, functional annotations, and how CF respiratory and environmental isolates are distributed on a core-genome phylogeny.

## Methods

### Samples

Our final dataset included 286 *P. aeruginosa* genome isolates downloaded from public genome repositories: 146 CF respiratory samples and 140 from environmental sources (Supplemental Table 1). ‘CF respiratory’ denotes isolates recovered from respiratory samples from pwCF; it is a sampling-source category and not a fixed evolutionary trait. Environmental isolates were associated with soil, water, or other non-human sources.^47–55^

Environmental identifiers were standardized to their base GCA accession numbers. When the same genome appeared across multiple filenames, we kept the standard ‘ASM…genomic.fna’ record, and an in vitro sample of PAO1 was treated only as a reference rather than an environmental observation. All 140 accepted environmental accessions matched the genome-summary table.

The CF respiratory cohort included 60 Danish patient–clone representatives from 33 patients, 82 American (Stanford) patient representatives, and four Italian patient representatives.^5,54,55^ For the Danish longitudinal samples, we kept only one observation for each patient-clone combination, but a patient could contribute more than one distinct clone-type. Accordingly, our unit of analysis is the sample unit defined in the archived dataset, rather than necessarily one unique patient per observation.

We obtained all bacterial genomes as preassembled genome sequences and did not perform raw-read trimming or de novo genome assembly. The original studies for the CF respiratory group and the public repositories for the environmental group supplied the assemblies used in our analysis.

The final prophage call tables determined which genomes were included in the analysis. We used a single canonical manifest to connect each genome ID with its source, cohort, patient or clone metadata when encoded in the identifier, bacterial genome length, GC content, and corresponding tip on the phylogenetic tree. For Figure 1, we retained genomes with length >5.9 Mbp and GC fraction >0.635, leaving140 CF respiratory and 139 environmental genomes. The reconciled phylogenetic subset contained 144 CF respiratory and 139 environmental genomes because two American samples and one environmental accession were absent from the supplied tree.

**Figure 1.**
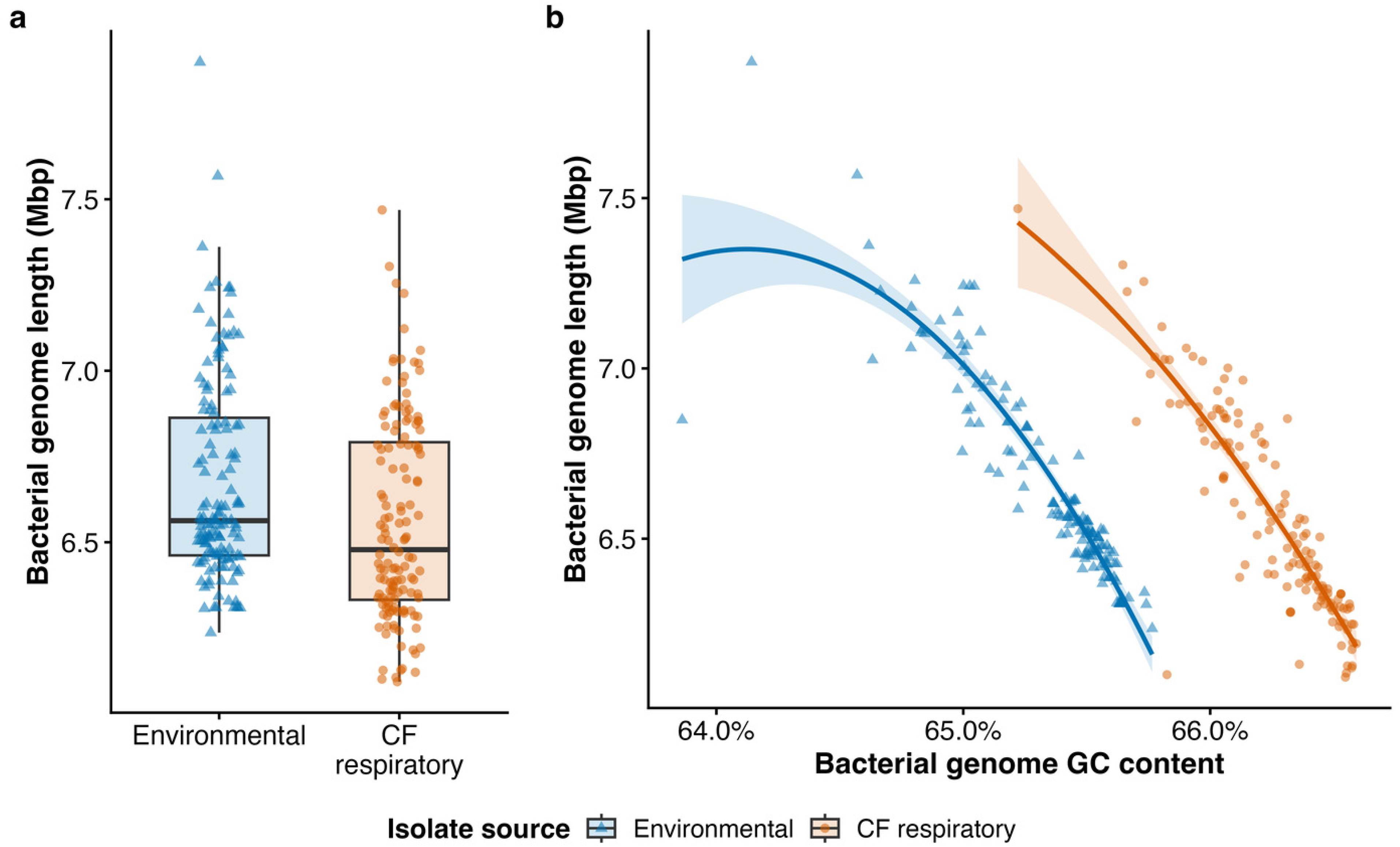
Bacterial genome length and GC content in CF respiratory and environmental *Pseudomonas aeruginosa* isolates. (a) Bacterial genome lengths for 140 CF respiratory and 139 environmental analysis units after the prespecified length (>5.9 Mbp) and GC (>0.635) thresholds. Points are analysis units; boxes show medians and interquartile ranges (IQRs), and whiskers extend to values within 1.5×IQR. (b) Relationship between bacterial genome GC content and bacterial genome length. Curves are source-specific quadratic-model predictions and shaded bands are 95% confidence intervals. Adding quadratic terms improved fit over the corresponding linear model (F=27.66, p=1.15×10^−11^); adjusted R^2^ was 0.79 for the linear model and 0.832 for the quadratic model.

### Mobile Genetic Element Database Construction

To identify proteins associated with *P. aeruginosa* MGEs, we constructed a custom database from NCBI nonredundant protein entries annotated with prophage-, phage-integrase-, structural-phage-, transposase-, insertion-sequence-, or recombinase-related terms and assigned to *P. aeruginosa* (accessed October 25, 2025). We grouped highly similar protein sequences using CD-HIT at 90% amino-acid identity. Assembly proteins were searched with BLASTp (E-value <10^−5^; minimum alignment coverage ≥80%).

### Genomic Analyses

We calculated each bacterial genome’s total length and GC content using the dplyr package in R. We defined GC content as the number of G and C bases divided by the total assembled genome length. We used no bespoke software. We then tested how genome length was related to GC content in CF respiratory versus environmental isolates using both linear and curved (quadratic) models. We compared the number of prophage-associated calls per genome using negative-binomial regression and an offset for log bacterial genome length. We also repeated the analysis without the supplied two-standard-deviation contig-length filter to check whether that filtering choice may have affected the result. Because each genome could contain multiple prophage-associated contigs, contig lengths were analyzed with a log-length linear mixed model containing a random intercept to account for multiple measurements from the same genome. Finally, we compared proportions of different MGE classes between groups using Wilcoxon rank-sum tests, and to account for the large size of our dataset in significance testing, we applied Benjamini–Hochberg correction across classes.

We classified phage-related sequences based on the markers found on each assembled contig. A contig containing both an identifiable integrase and at least one structural phage gene was labeled a high-confidence full-prophage call, while a contig with structural phage genes but without an identifiable integrase was labeled a fragment. Other marker combinations were assigned to their corresponding MGE categories.^58^ Importantly, then, the reported ‘length’ is the length of the entire labeled contig, and not the length of a boundary-resolved prophage genome.^70^

### Prophage sequence analysis

Prophage-associated proteins were annotated with Pharokka v1.9.0,^60^ including PHROG functional categories,^61^ VFDB virulence factors,^62^ and CARD antimicrobial resistance annotations.^63^ Category fractions were calculated as the number of proteins in a category divided by the total number of annotated proteins for the displayed unit. ‘Unknown function’ denotes proteins without a confident functional assignment; ‘other’ pools confident annotations that did not belong to the named categories. Because most proteins were of unknown function, that category is described separately in the results.

### Phylogenetic Analyses

We constructed a maximum-likelihood phylogenetic tree from the core-genome alignment. We used Parsnp in the Harvest^93^ suite to generate the core-genome alignment, using PAO1 as the reference genome and the -c option to include all input assemblies. Parsnp used its bundled FastTree2 implementation to infer an approximate maximum-likelihood core-genome SNP tree^94^. We imported the resulting Newick tree into R 4.4.1 with ape version 5.8-1, standardized the tip labels, retained one representation of each accession, and rooted the tree on PAO1. PAO1 was then removed before fitting two-state continuous-time Markov models. Equal-rates (ER) and all-rates-different (ARD) models were compared by AIC. Using the ER model and using a fixed seed, we ran 500 simulations of how isolates may have shifted between environmental and CF-respiratory states across the tree (stochastic character maps). We then calculated the difference between the environmental-to-CF-respiratory and CF-respiratory-to-environmental transitions using empirical 95% simulation intervals (Figure 4).

Analyses were conducted in R 4.4.1 with the packages ape^64^ and phytools.^65^

### Statistics

We performed all statistical analyses in R version 4.4.1. We used 2-sided Welch’s t-tests and Wilcoxon rank-sum tests to compare continuous outcomes between CF respiratory and environmental isolates.^59^ We used linear and quadratic regression models to evaluate the relationship between bacterial genome length and GC content and compared their fit with a nested-model *F* test. We analyzed prophage-associated contig counts with negative-binomial regression and included log bacterial genome length as an offset. We log-transformed prophage-labeled contig lengths and analyzed them with a linear mixed-effects model that included a random intercept for genome to account for multiple contigs from the same isolate.^57^ We compared isolate-level proportions of mobile genetic element classes with Wilcoxon rank-sum tests and corrected for our large dataset using the Benjamini-Hochberg method. We compared phylogenetic transition models using the Akaike information criterion and estimated uncertainty in transition counts from 500 stochastic character maps. We used 2-sided tests throughout, and considered p or adjusted p values below 0.05 significant.

## Results

### Bacterial genome size and GC content differ between CF respiratory and environmental isolates

We first asked how environmental and CF isolates of *P. aeruginosa* differed regarding their genome size and GC content. To this end, we calculated each bacterial genome’s total length and GC content.

We observed that CF respiratory bacterial genomes averaged 6.559±0.293 Mbp, compared with 6.675±0.296 Mbp for environmental genomes (Welch p=0.00122; Figure 1a). CF respiratory genomes also had higher average GC content (66.253%) than environmental genomes (65.320%; Welch p=7.74×10^−79^; Figure 1b). Genome length was negatively associated with GC content in both groups.

We then tested how genome length was related to GC content in CF respiratory versus environmental isolates using both linear and curved (quadratic) models, and compared the models to see which fit the data better with a nested F-test. Adding the source-specific quadratic terms improved fit over the corresponding linear model (F=27.66, p=1.15×10^−11^). Fitted quadratic curves are shown with 95% confidence intervals in Figure 1.^56^ Adjusted R^2^ increased from 0.79 for the linear model to 0.832 for the quadratic model. Figure 1b therefore shows quadratic fitted curves and 95% confidence bands, rather than describing nonlinearity from visual inspection alone.

These data indicate that CF respiratory isolate genomes were modestly smaller and substantially higher in GC content, and a quadratic model better represented the relationship between those variables.

### Prophages are shorter in CF compared to environmental isolates while the overall number of prophages does not differ between the groups

We next investigated the length and number of prophage sequences in these environmental versus CF respiratory infection isolates of *P. aeruginosa*.

Using the two-standard-deviation length filter, CF respiratory isolates had fewer high-confidence, full-prophage-labeled contigs on average than environmental genomes did: 4.95±3.43 high-confidence full-prophage-labeled contigs, compared with 8.51±4.79 in environmental isolates. After accounting for differences in bacterial genome size, CF respiratory genomes had about 42% fewer of these calls, as a negative-binomial model with a bacterial-genome-length offset estimated an incidence-rate ratio (IRR) of 0.583 (95% CI 0.506–0.672; p=8.12×10^−14^). This count contrast was not retained when all high-confidence full calls were analyzed: means were 10.70±3.30 and 11.04±4.49, respectively, and the IRR was 0.968 (95% CI 0.895–1.047; p=0.411).

Among the calls that passed the filter, full-prophage-labeled contigs were also shorter in CF respiratory isolates (mean 92.96±51.82 kbp; median 90.45 kbp) than in environmental isolates (mean 149.84±88.53 kbp; median 130.66 kbp) (Figure 2a). The mixed-effects model estimated that the geometric mean contig length was about 63% of that in environmental genomes (95% CI 0.584–0.677; p=8.15×10^−28^).

**Figure 2.**
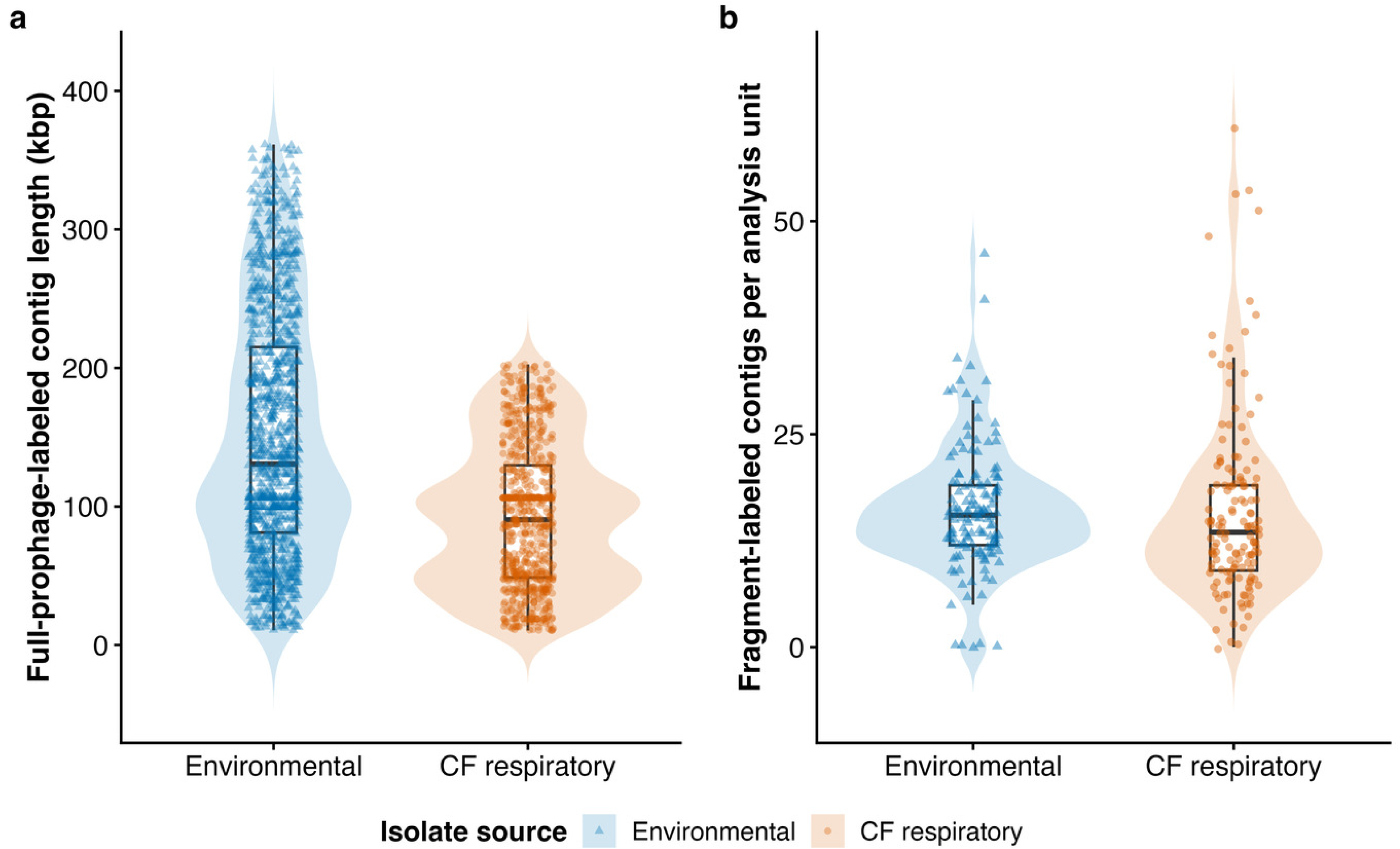
Prophage-labeled contig length and fragment counts by source. (a) Length distributions of high-confidence full-prophage-labeled contigs after the supplied two-standard-deviation length filter (CF respiratory n=722 calls; environmental n=1,192 calls). Points are contigs; boxes show medians and IQRs, and violins show the distributions. A log-length mixed model with an analysis-unit random intercept estimated a CF-respiratory/environmental geometric-mean ratio of 0.629 (95% CI 0.584–0.677). (b) Fragment-labeled contigs per analysis unit, including zeroes (CF respiratory n=146; environmental n=140). The genome-length-offset negative-binomial IRR was 0.976 (95% CI 0.862–1.105). Of note, contig lengths are not exact prophage-boundary lengths.

Fragment-labeled contig counts were similar between the two groups. CF respiratory genomes contained an average of 15.99±10.83 fragments, compared with 16.15±7.28 in environmental genomes (Figure 2b). The statistical model found no meaningful difference between them, with an IRR of 0.976 (95% CI 0.862–1.105; p=0.698).

From these data we conclude that that the filtered full-prophage-labeled contigs were shorter in CF respiratory infection than in environmental isolate genomes. In contrast, the number of prophages does not appear to differ between the groups.

### CF respiratory and environmental isolates are interspersed phylogenetically, without supported directional bias

We then investigated the phylogenic relationships between these isolates. After matching sample names to the tree, the phylogenetic analysis included 144 CF respiratory and 139 environmental study tips, plus PAO1 as the reference for rooting.

We observed that CF respiratory infection and environmental isolates were spread throughout the core-genome tree, rather than forming two source-specific groups (Figure 3). PAO1 was the only named reference in the supplied alignment.

**Figure 3.**
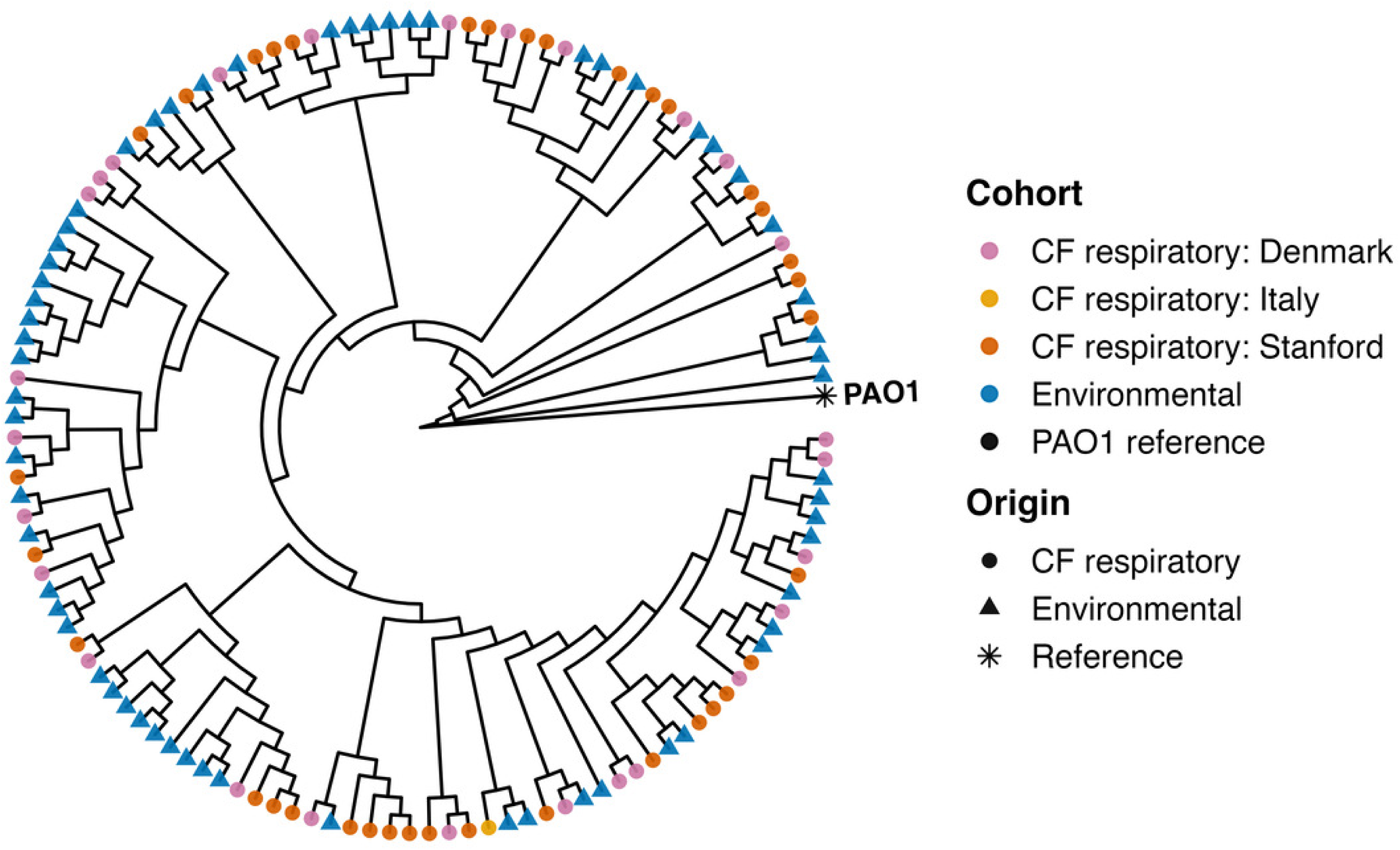
Core-genome phylogenetic distribution of CF respiratory and environmental *Pseudomonas aeruginosa* isolates. The reconciled tree contains 144 CF respiratory and 139 environmental study tips; PAO1 was retained as an outgroup landmark. For legibility, the circular cladogram shows a fixed, cohort-stratified 45% sample: CF respiratory Denmark n=27, Italy n=1, Stanford n=36; environmental n=62; and PAO1 n=1. Color identifies cohort and shape independently identifies origin. Branch lengths are unscaled in this display. Importantly, “CF respiratory” simply describes where a genome was sampled, and it is not an inherited bacterial trait.

The equal-rates model fit the data better than the ARD model did (AIC 424.11 versus 426.11; Akaike weights 0.731 and 0.269). Across 500 ER simulated mappings, we inferred an average of 71.06±15.03 environmental-to-CF respiratory transitions and an average of 84.45±13.06 CF respiratory-to-environmental transitions (Figure 4). The average difference between these directions was −13.39, and its empirical 95% simulation interval (−42 to 48) included zero. In other words, there was no statistical support for one direction of transition occurring more often than the other.

**Figure 4.**
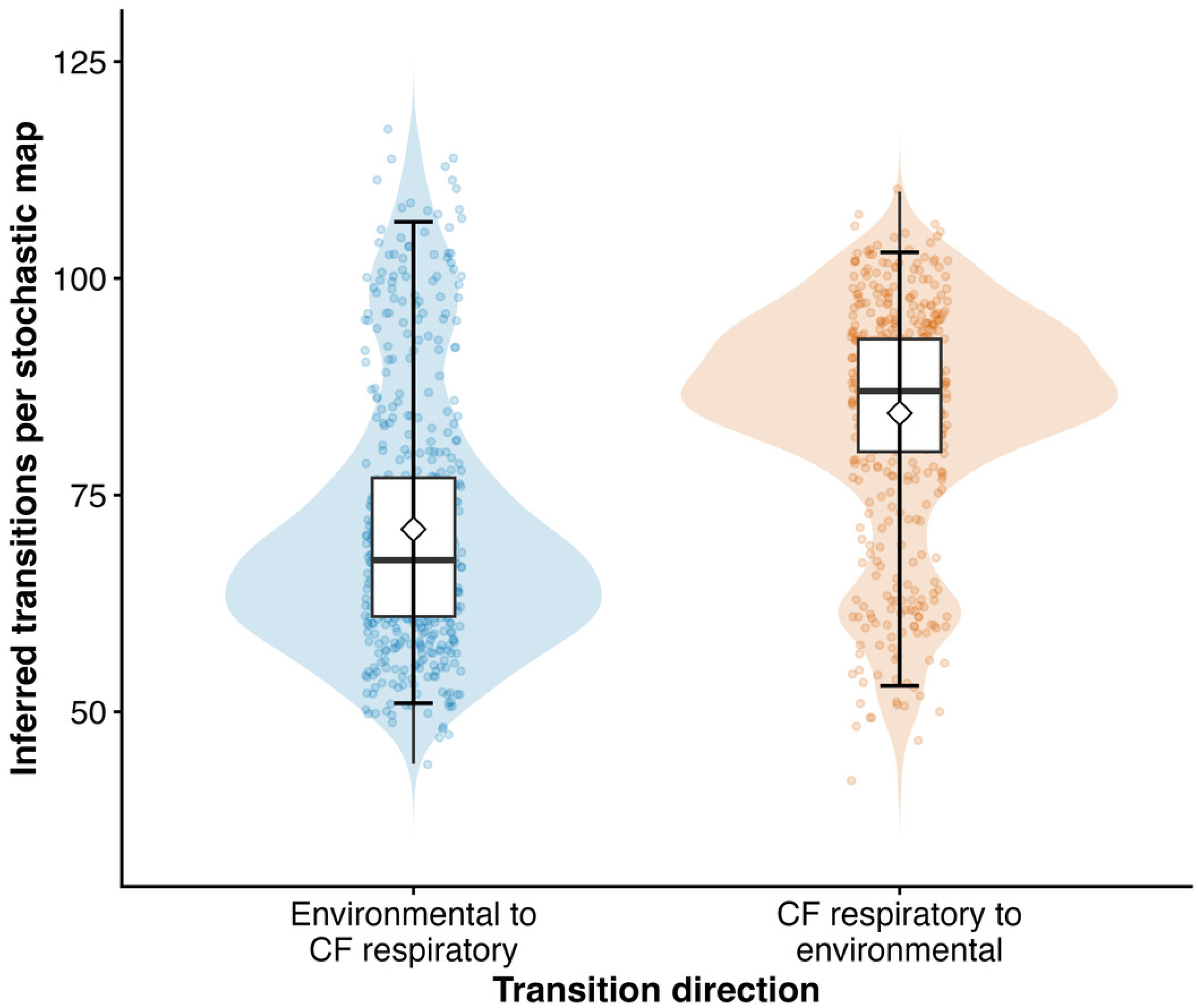
Source-state transitions inferred by stochastic character mapping. Counts were obtained from 500 equal-rates stochastic maps on the reconciled 283-tip study tree rooted with PAO1 before reference removal. Transparent points are map replicates; violins show distributions; boxes show medians and IQRs; diamonds show means; and error bars span empirical 95% simulation intervals. Environmental-to-CF-respiratory transitions averaged 71.06 (95% interval 51–106.5), and CF-respiratory-to-environmental transitions averaged 84.45 (95% interval 53–103). The directional-difference interval (−42 to 48) included zero. These data imply that pwCF acquire *P. aeruginosa* from environmental sources but also that clinical lineages are present in the environment.

Together, these phylogenic data show that CF respiratory and environmental isolates are repeatedly interspersed across *P. aeruginosa* diversity, but it does not support a claim that transitions preferentially occur in one ecological direction more frequently than the other. This implies that pwCF acquire *P. aeruginosa* from environmental sources but that isolates from clinical lineages are also present in the environment.

### The relative composition of identified MGE-labeled sequences are broadly similar between CF respiratory and environmental isolates

We then asked whether prophages, integrative elements, or transposons are significantly more abundant in either CF respiratory or environmental *Pseudomonas aeruginosa* isolates.

We observed that at the level of individual genomes, prophage-labeled sequence accounted for most of the mobile genetic element sequence in both groups: a mean of 81.51% of MGE-labeled sequence in CF respiratory isolates and 82.30% in environmental isolates (BH-adjusted Wilcoxon p=0.202; Figure 5). Integrative elements made up 17.62% and 16.59% respectively (BH-adjusted p=0.119) and 0.873% and 1.105% for transposons (BH-adjusted p=0.0507). These percentages refer only to the three displayed marker-defined classes and are not percentages of the bacterial genome.

**Figure 5.**
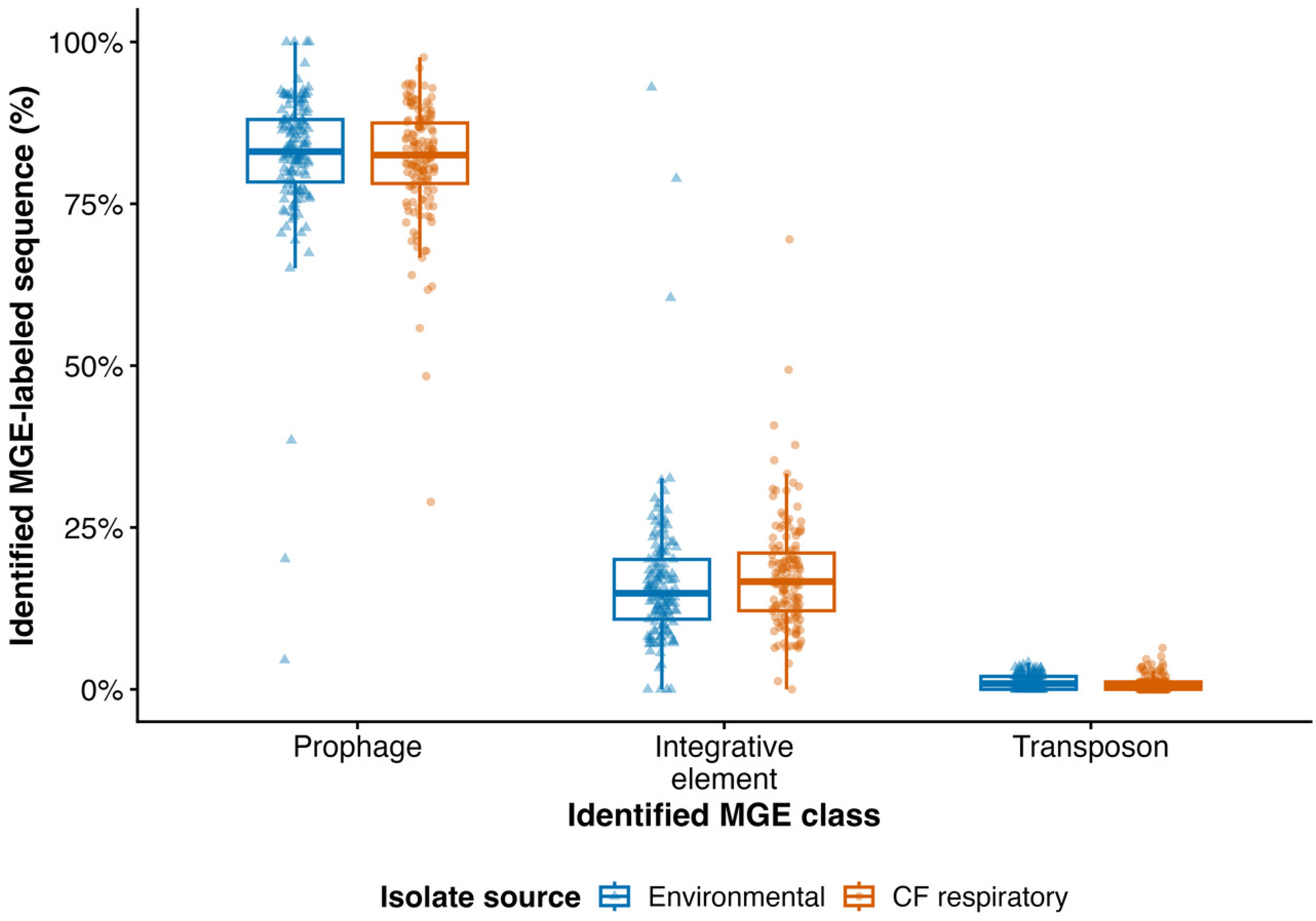
Analysis-unit composition of identified MGE-labeled sequence in CF respiratory and environmental *Pseudomonas aeruginosa* isolates. For each analysis unit, sequence assigned to prophage, integrative-element, or transposon classes was summed and expressed as a percentage of the total sequence assigned to those three classes. Points are analysis units; boxes show medians and IQRs. BH-adjusted Wilcoxon p values were 0.202, 0.119, and 0.0507, respectively. The denominator is identified MGE-labeled sequence, not total bacterial genome length. Plasmid was not a category in the supplied table. These data indicate that prophages, integrative elements, and transposons are not significantly more abundant in either CF respiratory or environmental *Pseudomonas aeruginosa* isolates.

These data indicate that prophages, integrative elements, and transposons are not significantly more abundant in either CF respiratory infection or environmental *Pseudomonas aeruginosa* isolates.

### Most prophage-associated genes have unknown function

We then investigated the function of prophage-associated genes in these isolates using functional annotations present in Pharokka v1.9.0,^60^ including PHROG functional categories,^61^ VFDB virulence factors,^62^ and CARD antimicrobial resistance annotations.^63^

We observed that the overall distribution of functional categories was also similar between CF respiratory and environmental genomes (Figure 6). Genes labeled ‘unknown function’ dominated both groups, at approximately 85% of annotated prophage-associated genes in the displayed distributions. This large uncharacterized fraction reflects the broader issue of with “viral dark matter” and limits functional interpretation.^72^ Of note, ‘Other’ denotes genes with confident functions that did not fit into one of the named categories, whereas ‘unknown function’ denotes the absence of a confident functional assignment. The archived antibiotic-resistance (CARD) and virulence-factor (VFDB) analyses did not show a source association after accounting for contig size.

**Figure 6.**
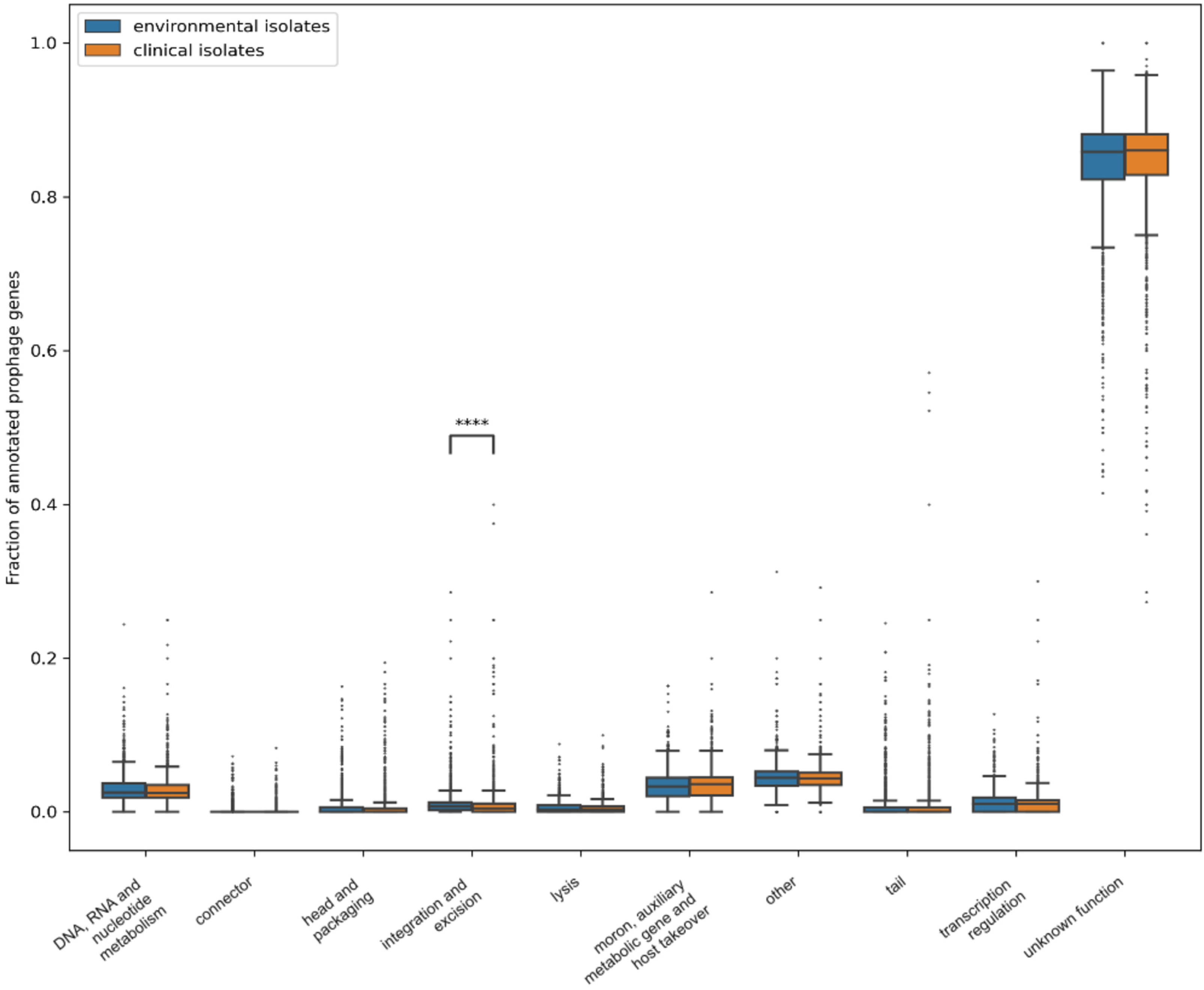
Functional categories of prophage-associated genes in environmental and CF respiratory isolates. Fractions are shown for named PHROG categories, an ‘other’ category pooling confident annotations outside the named categories, and ‘unknown function’ for genes without a confident assignment. Boxes show medians and IQRs; whiskers and points show the displayed spread and outliers. Unknown-function genes dominate both groups and are shown at the right. Integration and excision was the only category marked as different in the archived analysis.

Overall, “Integration and excision” was the only category that was different in this analysis. This suggests that genes involved in prophage mobilization in and out of the bacterial genome may be important to the adaptation to clinical environments. However, given the large number of ‘unknown function’ genes, it is difficult to know how much to make of this.

## Discussion

We have asked how bacterial genomes and prophages vary between *P. aeruginosa*isolates collected from CF respiratory infections versus environmental settings.

We find that CF respiratory isolate genomes were modestly smaller and higher in GC content. This combination of smaller genomes and higher GC content in the CF respiratory group is notable because extreme host association is often accompanied by genome reduction and, in many obligate pathogens and symbionts, lower GC content.^71^ *P. aeruginosa* is neither an obligate symbiont nor confined to one host niche, so the present contrast should not be treated as an exception to a universal rule. It may instead reflect differences in accessory content, lineage composition, or assembly characteristics that require boundary-resolved and lineage-aware testing. GC content can be shaped by many processes, including mutational bias, selection, codon usage, recombination, and horizontally acquired DNA, so these fitted relationships are best treated as evidence of a difference, rather than an explanation as to what caused that difference.

We find that filtered full-prophage-labeled contigs were shorter in CF respiratory isolates. The difference in the number of full-prophage calls result is less robust: the filtered analysis showed fewer full calls, but the unfiltered sensitivity analysis was null. Fragment counts were also null. Because each measurement is the length or count of a marker-labeled contig, these data cannot establish that environmental isolates carry more intact prophage genomes or that CF respiratory isolates have experienced greater prophage decay.

The source groups were interspersed on the core-genome tree, consistent with the recombining, weakly habitat-structured population described for *P. aeruginosa*.^66–69^ Although stochastic maps contained numerous changes between source labels, the directional-difference interval included zero. Consequently, the revised analysis does not support a preferred transition direction or prolonged residence in a ‘clinical’ state.

The predominance of genes with unknown function is itself an important result. Viral sequence space is dominated by poorly characterized genes, which constrains pathway-level interpretation and can obscure source-specific functions.^72^ The similar displayed distributions therefore indicate both broad functional similarity and limited annotation resolution, not evidence that the two prophage-associated gene pools are functionally identical.

This study has several limitations, starting with fragmented isolates and incomplete viral annotation. The Figure 1 thresholds and the supplied two-standard-deviation contig-length filter define analysis-specific subsets, and the latter changes the full-call abundance conclusion. The marker-based workflow does not provide exact prophage boundaries, host-flanking windows, plasmid calls, or a definitive lytic-versus-lysogenic classification. Phage-like contig in a bacterial assembly are not necessarily an integrated prophage.^70^ Lytic phage sequences can occur in bacterial isolates. Further, lysogenic phages can degrade and lose functionality, leaving behind degraded phage remnants like genomic scars.^95^ The archived inputs also do not support complete-prophage family assignment, prophage phylogeny, or localization of AMR and airway-adaptation genes to bounded prophages. Moreover, the large fraction of viral genes without confident functional assignments—the viral dark matter—limits functional and taxonomic inference.^72^

Future work should apply boundary-aware viral and plasmid detection to the original per-isolate isolates, validate integration sites and host flanks, compare prophage and host GC locally, cluster complete prophage genomes into families, and repeat phylogenetic analyses with named reference strains and complete lineage metadata. Those additions would test whether the present contig signatures reflect integrated prophages, assembly-associated lytic phages, plasmids, or other accessory elements and would clarify their functional relevance to CF airway biology. Moreover, longitudinal assessments of prophage contributions to evolutionary adaptation and experimental studies are needed, perhaps using established frameworks^91,92^.

In summary, these results support differences in bacterial genome composition and prophage-associated contig length between CF respiratory and environmental isolates. However, these studies are constrained by the limited information regarding prophages and much work remains to be done.

## Author contributions

RWT and PLB designed the research study. RWT, ZF, and JDP performed experiments and analyzed the data. RWT, ZF, PLB, and JDP contributed to the design of the figures. RWT, EBB, CEA, ZF, PLB, and JDP contributed to the writing of the manuscript.

## Acknowledgments

EBB was supported by the NIH/National Heart, Lung, and Blood Institute (grant 1K23HL169902-01) and the Cystic Fibrosis Foundation (grants BURGEN23G0 and BURGEN24A0-KB). P.L.B. discloses support from National Institutes of Health grant R01 HL148184-01; R01 AI12492093; R01 DC019965, K24 AI166718, P01AI1960471, and R01 EB038154 and support from the CFF and the CFRI. The contents are those of the authors and do not necessarily represent the views of the funding agencies.

